# Optimizing Electroporation Parameters for Efficient Delivery of Large Molecules into Pig Zygotes Using Fluorescent Dextrans from 3 to 2000 kDA

**DOI:** 10.1101/2024.08.26.609025

**Authors:** Juan Pablo Fernández, Paul Kielau, Petra Hassel, Wilfried A. Kues

## Abstract

Electroporation has revolutionized gene transfer and gene editing, enabling efficient delivery of molecules into embryos, with significant implications for developmental biology and biomedical research. This study aimed to optimize electroporation parameters for enhancing the delivery of large molecules into pig zygotes. We investigated the effects of fluorescence-coupled dextran reporters (FDs) of sizes ranging from 3 to 2000 kiloDalton (kDA) along with the impact of poring and transfer polarity settings during electroporation, on molecule permeability. Additionally, we assessed the influence of voltage and the number of poring pulses on the delivery of 2000 kDa FDs and examined the permeability of pre-IVF embryos and zona pellucida-weakened post-IVF embryos to this FD.

Our findings highlighted size-dependent effects on FD uptake, with reversing poring polarity increasing the influx of small molecules (3 kDa FDs). The delivery of 2000 kDa FDs was not influenced by increased poring number but it was significantly influenced by voltage, reaching its optimum at 40 V. Electroporation in pre-IVF embryos did not show significant variation across different voltages. However, voltages higher than 20 V negatively affected blastocyst development rates. Zona-weakening did not improve permeability for the 2000 kDa FD.

This study offers valuable insights into refining electroporation techniques for delivering large molecules into pig zygotes and highlights the relevance of commercial fluorescence-coupled dextrans as useful tools for exploring permeability dynamics in electroporated zygotes.

## Introduction

Developing new methods for delivering genetic material into zygotes that are less labor-intensive, faster, and easier to handle than microinjection techniques is highly desirable(1). The application of electroporation in genetic manipulation has revolutionized the field of molecular biology by enabling efficient delivery of molecules into cells(2, 3). Particularly in reproductive biotechnology, electroporation has emerged as a powerful tool for introducing genetic material into zygotes, offering unparalleled opportunities for gene transfer and genome editing(4). However, while the efficacy of electroporation in delivering molecules into cells is well-established, the permeability dynamics of different-sized molecules in zygotes remain poorly understood. Electroporation is particularly effective for introducing ribonucleoproteins (RNPs), RNAs, and short DNA sequences into zygotes(4). However, these molecules are significantly smaller than plasmids, which are essential for the production of transgenic animals and can range from 1,000 to over 50,000 base pairs (bp). With each base pair weighing about 0.65 kilodaltons (kDa), plasmids are much larger than the molecules typically used in electroporation.

In the context of using pigs as a biomedical tool for research, DNA transfection is crucial(5). In the field of xenotransplantation, for instance, it is necessary to delete xenoantigens, but also introduce human genes into pig embryos(6). This dual modification enhances the compatibility and acceptance of pig organs in human recipients(7).

Embryo electroporation techniques aim to maximize gene editing efficiency while minimizing damage to embryos. Researchers have examined factors such as the number, duration, and voltage intensity of electroporation pulses to enhance molecule entry into zygotes without adversely affecting blastocyst formation rates(8, 9).

The TAKE method developed by NepaGene distinguishes itself from other electroporation devices with its dual-pulse mode, which is particularly useful to ensure high transfection efficiency(2, 10). Furthermore, it allows for independent configuration of poring and transfer pulse polarity within a single electroporation run, enabling a wide variety of experimental combinations(11).

In addition to optimizing electroporation settings, treating embryos with pronase, a mixture of proteolytic enzymes from the bacterium *Streptomyces griseus*, to weaken or remove the zona pellucida has been reported to improve transfection efficiency(12, 13). This enzymatic treatment might increase the permeability of the zona pellucida, facilitating greater molecule entry. Moreover, electroporating early-stage or unfertilized embryos has been found to be more effective for gene editing than later-stage embryos(14, 15), potentially due to the rapid delivery of Cas9 before cleavage. However, the morphological characteristics and permeability of early-stage or unfertilized embryos may also play a role.

Commercial fluorescence-coupled dextrans (FDs) are accessible and ready-to-use molecules that can be detected just 2 hours after electroporation to evaluate molecule entry. This is a significant advantage over the lengthy screening process of gene editing analysis, which requires PCR, endonuclease assays, or further sequencing steps of the resulting blastocysts. Moreover, these globular fluorescent reporters are available in specific sizes, serving as reliable references for other molecules.

In this study, we aim to elucidate how the fluorescence intensity of variously sized dextrans can serve as a proxy for understanding the permeability and delivery efficiency in pig zygotes for specific electroporation approaches. This knowledge will help optimize electroporation techniques for the successful delivery of larger molecules, enhancing the potential for gene-editing applications in developmental biology and other fields.

## Project Description

This project aimed to optimize electroporation parameters for delivering large molecules into pig zygotes. We used dextran fluorescent dextrans (FDs) of varying sizes to visually assess the permeability of pig zygotes. The project consists of three studies, each focusing on different aspects of the electroporation process.

### Study 1: Impact of FD Size and Dual Polarity Setting in Molecule Permeability via Electroporation

This study assessed the influence of poring and transfer pulse polarity settings on molecule permeability and blastocyst formation rate (BR) during electroporation. Pooled fluorescent dextrans (FDs) of 3, 70, and 500 kDa were used to measure fluorescence intensity and BR across four possible combinations of poring (P) and transfer (T) pulse polarities. These parameters may potentially affect how efficiently molecules are introduced into the cells. The four dual-polarity (P/T) combinations tested were:

P+/− T+/− (alternating polarity for both poring and transfer pulses)
P+/− T+/− (alternating polarity for poring and stable polarity for transfer pulses)
P+ T+/− (stable polarity for poring and alternating polarity for transfer pulses)
P+ T+ (stable polarity for both pulse and transfer pulses)

### Study 2: Impact of Voltage and Number of Poring Pulses on the Electroporation of 2000 kDa FDs in Post-IVF Porcine Embryos

The optimal (P/T) polarity combination resulting from study 1 was used to individually electroporate a larger FD (2000 kDA) into porcine zygotes. First, we increased poring pulse frequency to five and six pulses, while maintaining a constant voltage of 30 V. Conversely, we also maintained a four-pulse frequency constant, while gradually increasing voltage levels from 20 to 50 volts.

### Study 3: Impact of Electroporation in Pre-IVF and Zona Pellucida-weakened Post-IVF Embryos in Molecule Permeability

This study evaluated the efficiency in molecule permeability during electroporation when performed on oocytes prior to fertilization, aiming to understand its impact on molecule entry and early-stage embryo development. Additionally, we also explored how the weakening of the zona pellucida affects these two factors.

## Materials and Methods

### Ethics Declaration

The experiments were performed using ovaries from slaughterhouse material

### Ovary collection and maturation

Porcine ovaries were obtained from a local slaughterhouse and underwent initial cleansing in a 0.9% NaCl (Roth #3957.2, Karlsruhe, Germany) solution supplemented with 0.06 g/l penicillin (AppliChem #A1837, Darmstadt, Germany) and 0.131 g/l streptomycin (AppliChem #A1852,0250) before oocyte retrieval. Aspiration of the oocytes was carried out using a house vacuum connected to a Falcon tube. Subsequently, the collected oocytes were washed in porcine X Medium (PXM) (16). Adequate cumulus-oocyte-complexes (COCs) were defined by uniform cytoplasm and several layers of cumulus cells encompassing the zona pellucida. For maturation, the oocytes were transferred into FLI maturation medium(16). Each collection involved the incubation of a minimum of 300 COCs for 40-44 hours at 38°C in a humidified air environment with 5% CO2.

### Extraction, Freezing and Thawing of Porcine Semen

Sperm was obtained using the hand-gloved method with a phantom, filtered through gauze, and diluted 1:1 with prewarmed (38°C) Androhep Plus extender (Minitube, Tiefenbach, Germany). The diluted semen was transferred to the laboratory and kept in 50 mL centrifugation tubes at room temperature for 60 minutes. The semen was then cooled to 15°C over 90 minutes in an incubator. Next, the tubes were centrifuged at 800 x g and 15°C for 10 minutes to reduce seminal plasma. The supernatant was discarded and replaced with a chilling extender (80% 322 mM lactose solution + 20% egg yolk) equilibrated at 15°C. The sperm concentration was adjusted to 1.8 billion spermatozoa/mL using a NucleoCounter (ChemoMetec A/S, Allerod, Denmark). The semen was cooled to 5°C over 90 minutes and maintained at 4°C for 30 minutes.

All further steps were conducted at 4°C. Chilled semen was mixed 2:1 with a freezing extender (92.5 mL chilling extender, 1.5 g Orvus ES paste (OEP, Minitube), 6.0 g glycerol) to achieve a final concentration of 1.2 billion sperm/mL and 1.74% glycerol. Straws (0.25 mL; Minitube) were filled and sealed with an MPP Uno machine (Minitube). Freezing was performed in a Styrofoam box with liquid nitrogen. The samples were placed on metal racks 4 cm above the liquid nitrogen for 20 minutes, then plunged into liquid nitrogen and stored in cryo containers. Straws with frozen semen were plunged in a water bath at 37°C for 17 seconds. Then, the outer surface of the straws was carefully dried, the straws were cut open, and the content was expelled into a 3 ml Androhep-filled 15 ml Falcon tube. Centrifugation took place at 867 x g for three minutes at 30°C, and the pellet was washed again with 3 ml Androhep before being resuspended in 0.5 ml Fert-Talp(17)

### In-vitro-Fertilization (IVF)

The matured oocytes were coincubated for 5 hours with frozen-thawed ejaculated spermatozoa (1 × 10^6 cells/mL) in porcine fertilization medium (17), at a ratio of 60 spermatozoa to one oocyte. Then, the embryos were cultured in PZM in a humidified incubator at 39°C with 5% CO2 until the cytoplasmic maturation, pronuclear formation, and the introduction of TMR-Ds. When electroporation was performed before IVF, oocytes were subjected to electroporation 42 hours after incubation in FLI maturation medium and transferred to fertilization medium before coincubation with frozen-thawed spermatozoa occurred.

### Zona weakening of fertilized zygotes

As part of study 3, the zona pellucida of porcine zygotes was weakened using pronase prior to electroporation. Zygotes were treated with Pronase (Sigma Aldrich, Germany) diluted in TL Hepes 296 plus Ca to a final concentration of 1 mg/ml for 10 seconds. Following this, they were washed with TL Hepes containing 20% Newborn Calf Serum (Gibco) to halt the pronase activity and subsequently transferred to PZM. The zona pellucida-weakened zygotes were then incubated at 39°C in a humidified incubator with a gas mixture of 5% CO2, 5% O2, and 90% N2 for 1 hour before electroporation.

### Fluorescent dextrans and electroporation

For FD electroporation, the following lysin fixable dextran compounds ranging from 3 kDA to 2000 kDA were bought from Thermofisher (Waltham, USA):

- Dextran, Cascade Blue (3,000 MW, D7132)
- Dextran, Tetramethylrhodamine (70,000 MW, D1818)
- Dextran, Fluorescein (500,000 MW, D7136)
- Dextran, Tetramethylrhodamine (2,000,000 MW, D7139)

Dextrans were diluted in TL-Hepes Ca free to a final concentration of 2 mg/mL. Before electroporation, dextrans were vortexed, briefly heated at 40°C, and centrifuged at 12,000 × g for 5 minutes to remove insoluble particles. For Study 1, the 3 kDa, 70 kDa, and 500 kDa FDs were first electroporated individually to establish calibration settings for each fluorescent conjugate. For P/T polarity orientation analysis, these three dextrans were pooled at equal volumes for electroporation. In studies 2 and 3, further electroporation of a 2000 kDa FD was performed in pre-IVF oocytes and zygotes.

For the electroporation process, groups of 25 embryos each were washed three times in Opti-MEM before being placed in a chamber (CUY505P5, Nepa Gene; Chiba, Japan) containing 20 μL of the dextran solution. Electroporation was performed in a Nepa 21 electroporator (Nepa Gene, Chiba, Japan). Parameters were set as outlined in Table 1. After electroporation, zygotes were removed from the chamber using a glass capillary pipette, followed by a triple wash in TL-HEPES and an additional triple wash in PZM or fertilization medium (study 3) for subsequent culture.

**Table 1.** Parameters utilized for embryo electroporation.

| Parameters |  |  |  |  |  |  |  |  |  |  |  |
| --- | --- | --- | --- | --- | --- | --- | --- | --- | --- | --- | --- |
| Poring Pulse |  |  |  |  |  | Transfer Pulse |  |  |  |  |  |
| V<sup>2,3</sup> | Length (ms) | Interval (ms) | No.<sup>2</sup> | D.Rate (%) | Polarity<sup>1</sup> | V | Length (ms) | Interval (ms) | No. | D.Rate (%) | Polarity<sup>1</sup> |
| 20/30/<br>40/50 | 3.5 | 50 | 4/5/6 | 10 | +/-<br>+ | 5 | 50 | 50 | 5 | 40 | +/-<br>+ |

Parameters with superscripts denote variable settings applied in Study 1 (<sup>1</sup>), Study 2 (<sup>2</sup>) and Study 3 (<sup>3</sup>). V: Voltage in volts, D: Decay.

### Evaluation of fluorescence intensity in embryos post TMR-Dextran delivery

The fluorescence intensity emitted by pig embryos or oocytes following the delivery of FDs via electroporation was used as an indicator of molecular uptake. This intensity was documented two hours after electroporation or after in vitro fertilization (IVF) when electroporation was performed on oocytes. Four-well rectangular dishes (Thermo Fisher) containing the zygotes were placed under a confocal laser scanning microscope (Leica Stellaris 5 – Leica Microsystems, Wetzlar, Germany) and the corresponding software LAS X Life was utilized for analysis. The channels chosen for our imaging recordings were Cascade Blue for 3 kDa dextran, Alexa 488 for 500 kDa dextran, and Texas Red for 70 kDa and 2000 kDa dextrans.

First, zygotes that had not been electroporated underwent microscopical recordings to discard any autofluorescence signal that could interfere with the analysis and for background fluorescence analysis. To ensure that no crosstalking was present during our recordings, a spectral calibration using single-stained zygotes for each fluorophore was performed to ensure that the emission spectra did not overlap significantly. The resulting setting was utilized to acquire sequential images of zygotes electroporated with pooled dextrans.

Overlays were subsequently analysed by ImageJ (version 1.48) (18) to quantify fluorescence values. This quantification involved subtracting the fluorescence intensity emitted from zygotes from the fluorescence of the background as well as the background of non-electroporated embryos.

The linear regression model utilized to evaluate the effects of P/T polarity combinations on the fluorescence intensity resulting from zygotes is indicated as follow:

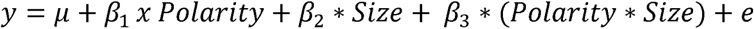

where *y* represents the fluorescence intensity, *µ* the overall intercept, *Polarity* is the effect of the combination of P/T polarity settings, and *Size* represents the fluorescence-coupled dextran size. *β*_1_, *β*_2_ and, *β*_3_ are regression coefficients, and *e* represents the residual error. For the effect of polarity in specific dextran sizes, the linear model follows the formula:

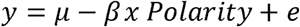

The effects in the model were tested for significance using an Analysis of Variance (ANOVA) on the linear model in R version 4.3.0 (19). For studies 2 and 3, fluorescence intensity was compared between different pulse number or voltage level groups by ANOVA followed by paired t-test.

## Results

### Impact of FD Size and Dual Polarity Setting in Molecule Permeability via Electroporation

When electroporating 3 kDA (blue), 70 kDA (red) and 500 kDA (green) FDs individually to calibrate intensity settings for each channel to avoid crosstalk between channels, the zona pellucida of embryos consistently emitted red fluorescence, regardless of the FD that was utilized (Figure 1A). This issue could not be avoided and fluorescence was only measured inside the zygotes to ensure accurate measurements. In contrast, electroporation in pre-IVF oocytes was performed this procedure eliminated the red autofluorescence in the zona pellucida (Figure 1B). When pooling 3, 70 and 500 kDA FD sizes for electroporation in zygotes, regression analysis indicated a significant effect of FD size on fluorescence intensity (P < 0.001), with smaller sizes yielding higher fluorescence values.

**Figure 1:**
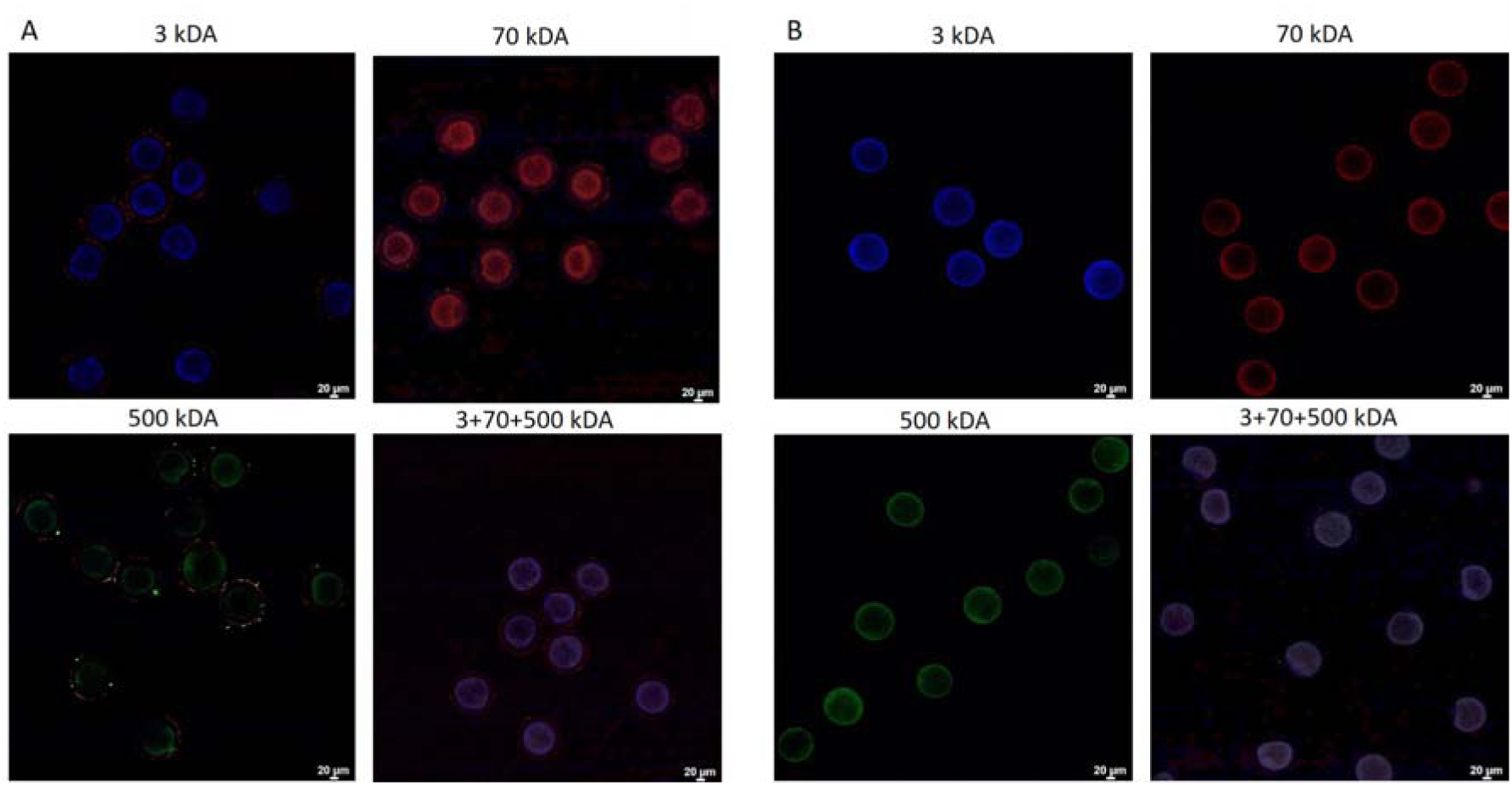
Electroporation of individual and pooled FDs in post- and pre-IVF oocytes. Fluorescence recordings following the electroporation of 3 kDa, 70 kDa, 500 kDa, and pooled FDs (3+70+500 kDA) in (A) IVF zygotes and (B) pre-IVF zygotes

P/T polarity combination settings did not significantly affect fluorescence values across FD sizes. However, an interaction between polarity and FD sizes was identified (P < 0.05). Specifically, the effect of polarity in 3 kDa FD was significant (P < 0.01), with the polarity combination P+ T+/− inducing the highest fluorescence intensity. This effect was not observed in 70 kDa and 500 kDa FDs (Figure 2). In relation to blastocyst rate, ANOVA showed no significant differences across the poring and transfer polarity combination settings Table 2.

**Figure 2:**
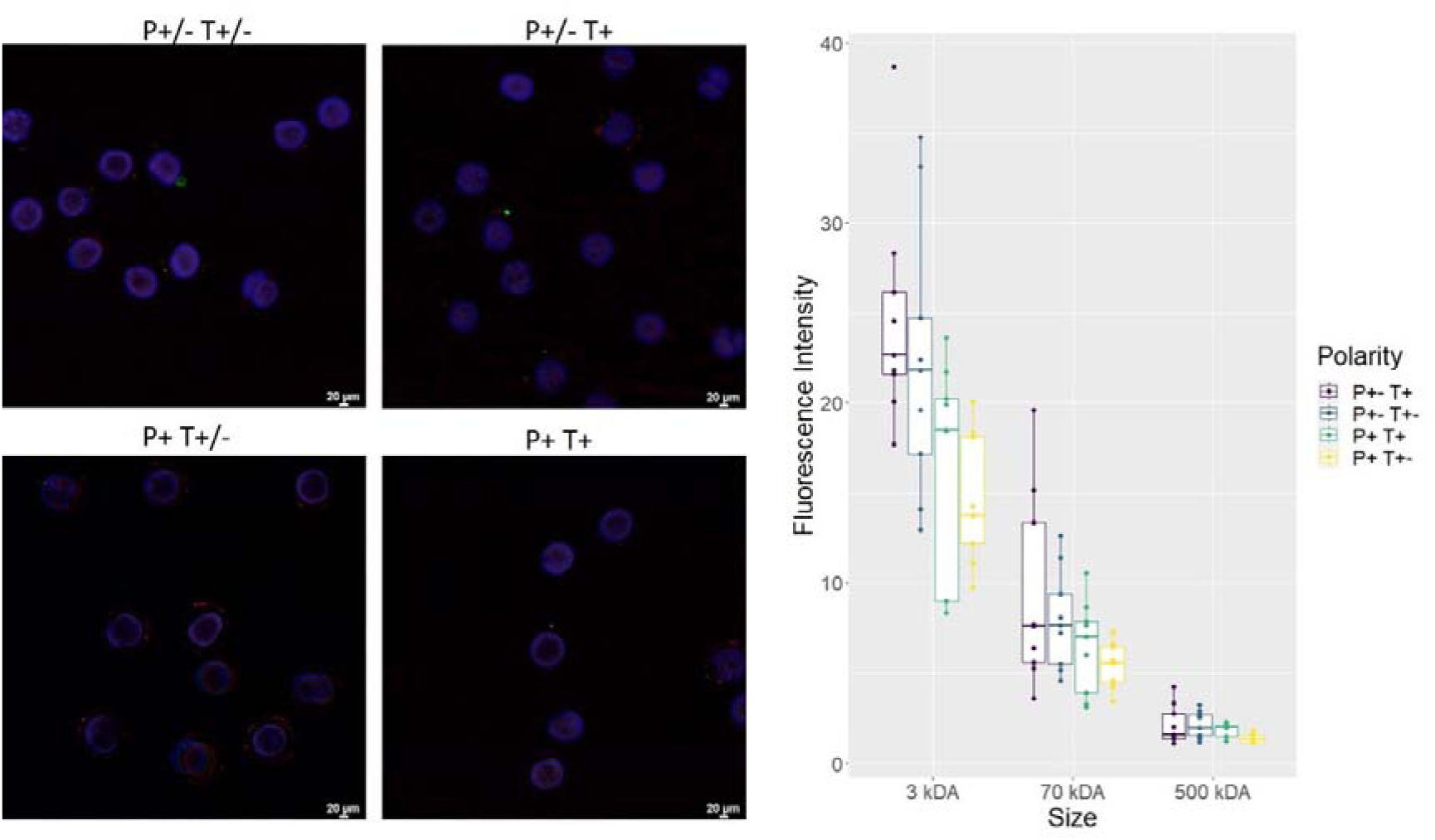
Fluorescence intensity across different dual-polarity setting combinations in the electroporation of pooled FDs in IVF embryos. (Left) Normalized images of pooled FDs delivered to embryos under specific dual-polarity combinations. (Right) Quantification of fluorescence values 2 hours post-electroporation reveals a significant effect of FD size on fluorescence intensity (P<0.001). An interaction between dual-polarity settings and fluorescence intensity is also demonstrated (P<0.05), with polarity combination settings showing a significant effect in the delivery of 3 kDA FDs (P<0.01).

**Table 2.**
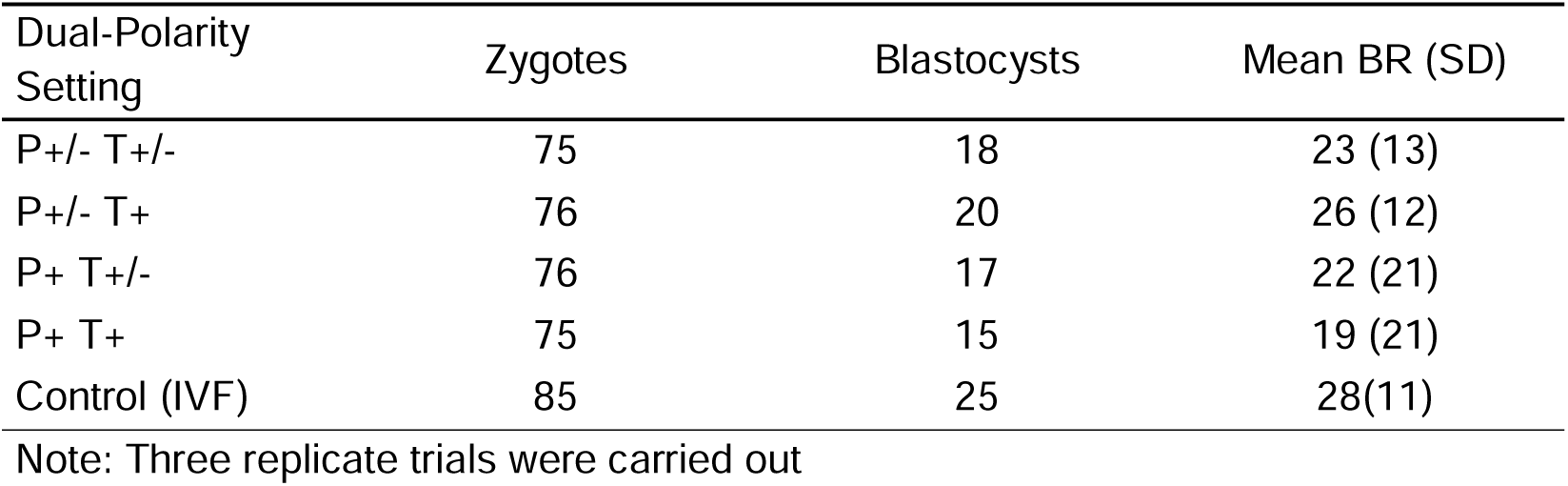
Analysis of BR for different polarity setting combinations in the electroporation of IVF embryos. No statistical evidence of differences in the mean BR was found through t-test.

| Dual-Polarity Setting | Zygotes | Blastocysts | Mean BR (SD) |
| --- | --- | --- | --- |
| P+/- T+/- | 75 | 18 | 23 (13) |
| P+/- T+ | 76 | 20 | 26 (12) |
| P+ T+/- | 76 | 17 | 22 (21) |
| P+ T+ | 75 | 15 | 19 (21) |
| Control (IVF) | 85 | 25 | 28(11) |

Note: Three replicate trials were carried out

### Impact of Voltage and Number of Poring Pulses on the Electroporation of 2000 kDa FD in Porcine Zygotes

In this study, a 2000 kDa FD was individually electroporated into IVF zygotes. First, the number of poring pulses was increased from four poring pulses to five and six pulses, while maintaining a constant poring voltage. Increasing the pulse number to 5 or 6 did not significantly improve fluorescence intensity (Suppl. Figure 1).

Next, the voltage levels were gradually increased while maintaining a constant number of 4 poring pulses. At 40 volts, a significantly stronger fluorescence intensity was observed compared to the other voltage levels (20 volts, P<0.001; 30 volts, P<0.001; or 50 volts, P<0.05). However, it was also observed that not all zygotes intended for 2000 kDA FD electroporation were successfully transfected at both 40 volts and 50 volts, with the number of successfully targeted embryos decreasing as the voltage increased (Figure 3).

**Figure 3:**
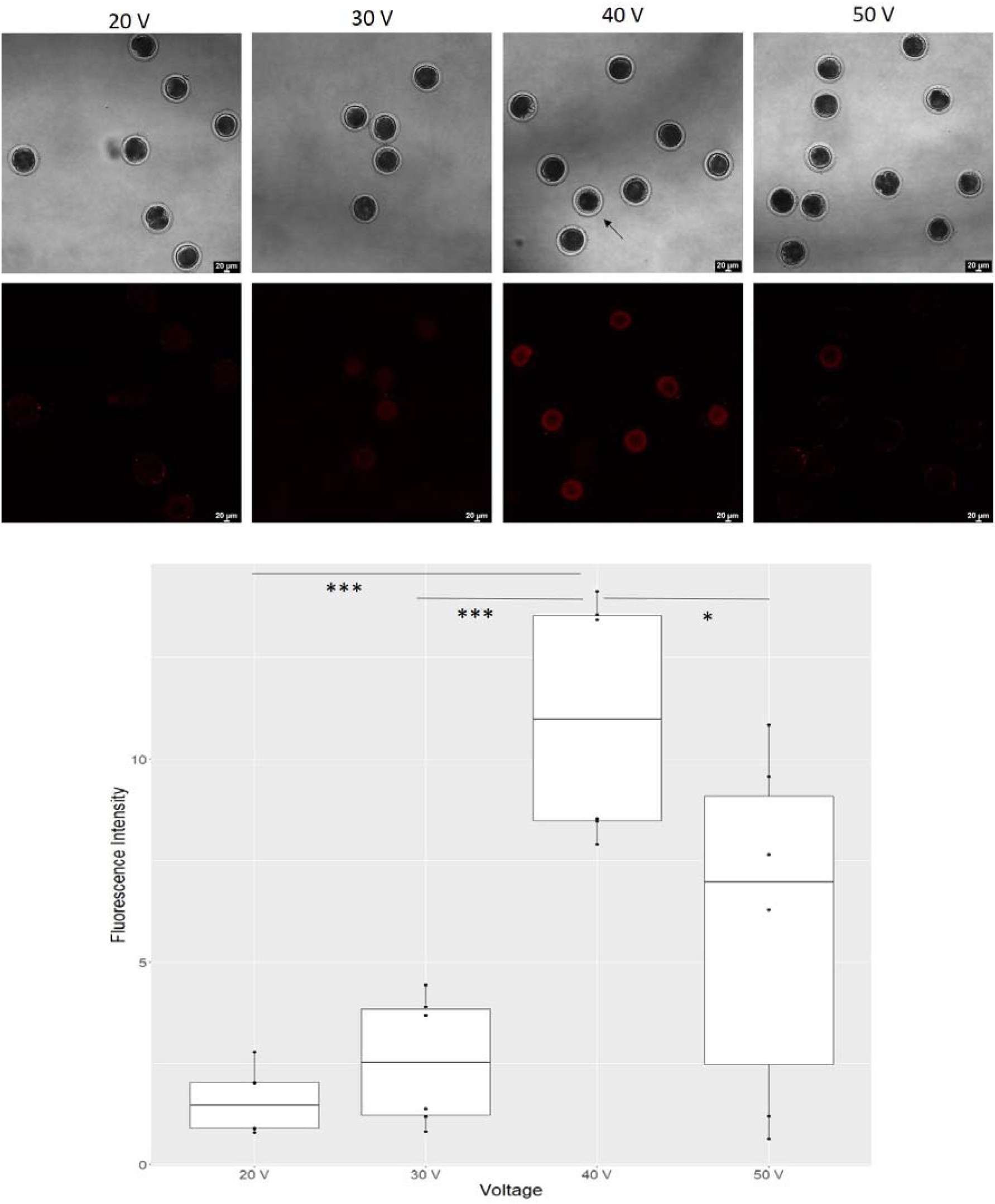
Electroporation of 2000 kDa FDs in IVF Zygotes with Incrementally Increasing Voltages. Higher voltages increased fluorescence intensity of 2000 kDa FDs, but also decreased transfection rate in zygotes. At 40 V, fluorescence intensity reached its average peak level. *** (P<0.001), * (P<0.05). The arrow indicates a zygote that failed to be transfected at 40 V.

**Figure 4:**
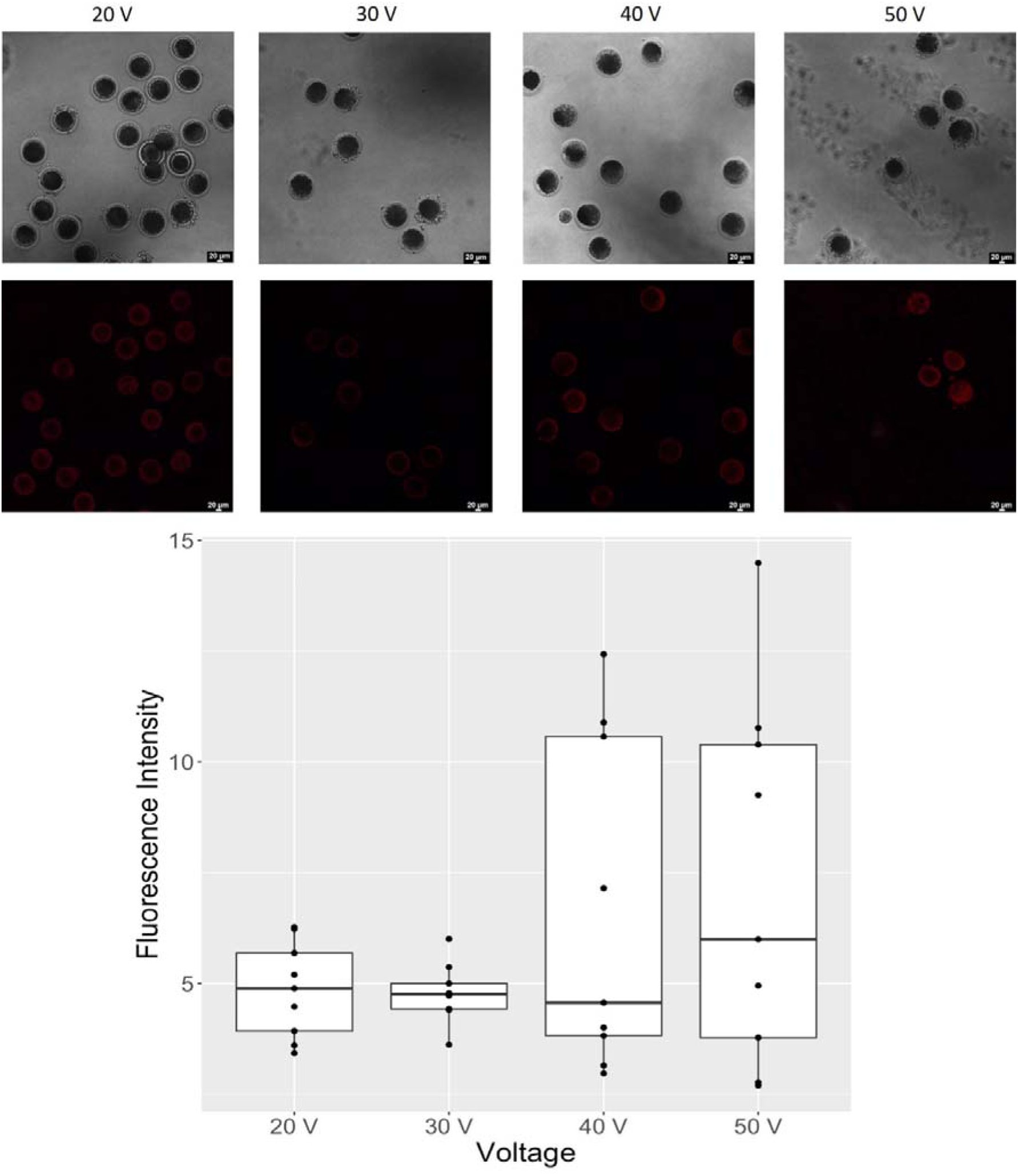
Electroporation of 2000 kDa FDs in pre-IVF oocytes with incrementally increasing voltages. No significant differences between fluorescence intensity values for the applied voltages

### Impact of Electroporation in Pre-IVF and Zona Pellucida-weakened Post-IVF Embryos in Molecule Permeability

In this study, pre-IVF electroporation of zygotes was performed to anaylse the permeability of zygotes before IVF carried out. The voltage-dependent electroporation of 2000 kDa FD in pre-IVF oocytes displayed a similar tendency to that of post-IVF electroporation, with oocyte fluorescence intensity increasing in relation to voltage increase and the number of fluorescent zygotes also decreasing.

Despite this trend, there were no significant differences in fluorescence intensity between different voltage levels. Blastocyst formation rate, however, was significantly lower in groups subjected to voltages above 20V compared to the IVF control group. To determine if voltage levels affected the blastocyst rate (BR), a 0-volt control was included. Electroporation at 0 volts did not differ from the IVF control group in terms of blastocyst rate development (Table 3). Although there was no statistical difference between the 30-volt and 40-volt groups compared to the 0-volt group, there was a decrease of approximately 22 percentage points in blastocyst formation rates.

**Table 3:**
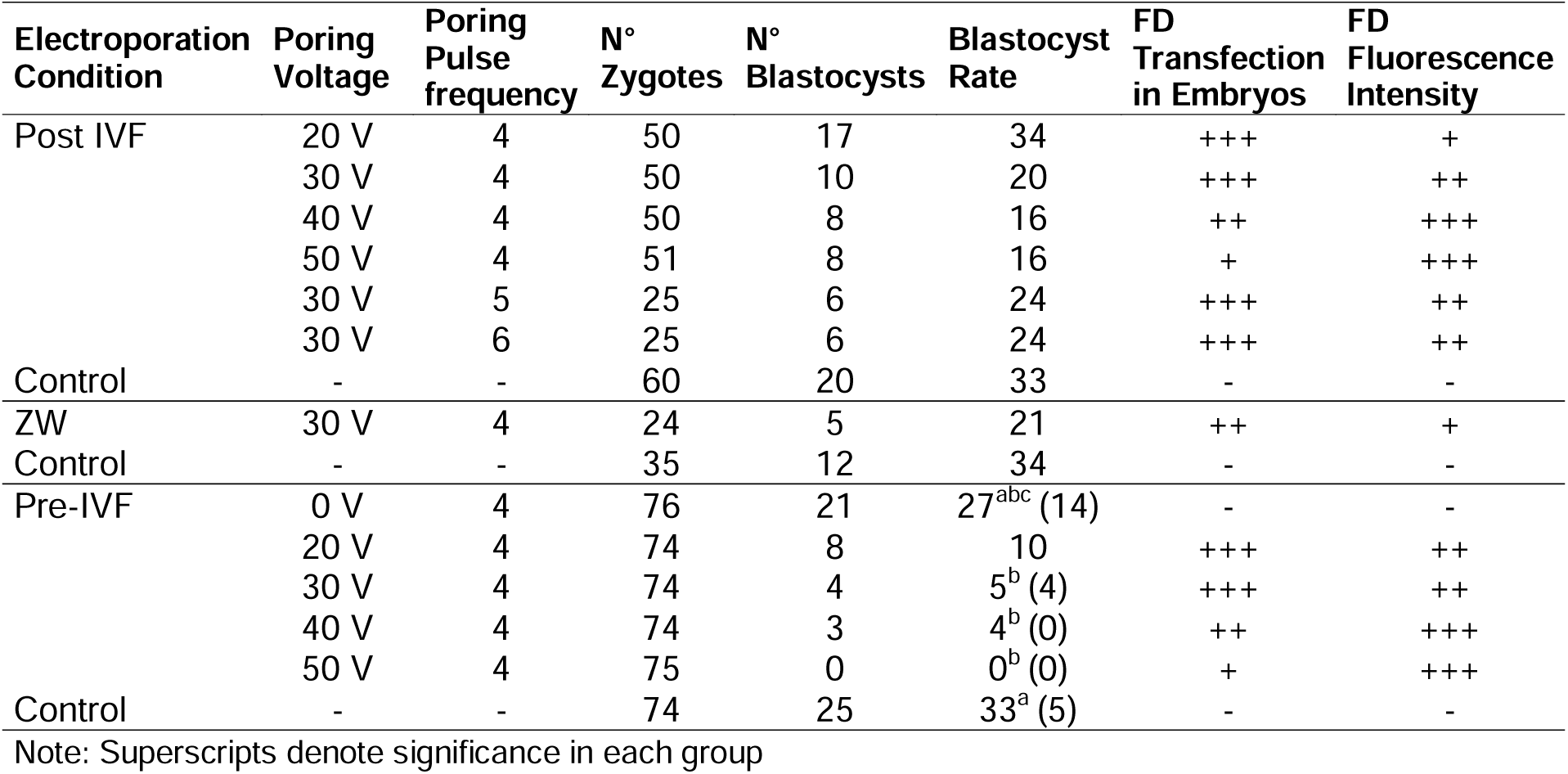
Summary of electroporation of 2000 kDa FDs in pre- and post-IVF embryos for BR and fluorescent characteristics under different conditions.

Regarding zona-weakened (ZW) zygotes, pronase activity was able to visually thin their zona pellucida. However, at 30 volts and 4 pulses, this treatment did not facilitate the entry of 2000 kDa FDs into embryos (Figure 5). BR was not significantly different between zona-weakened zygotes and controls (Table 3).

**Figure 5:**
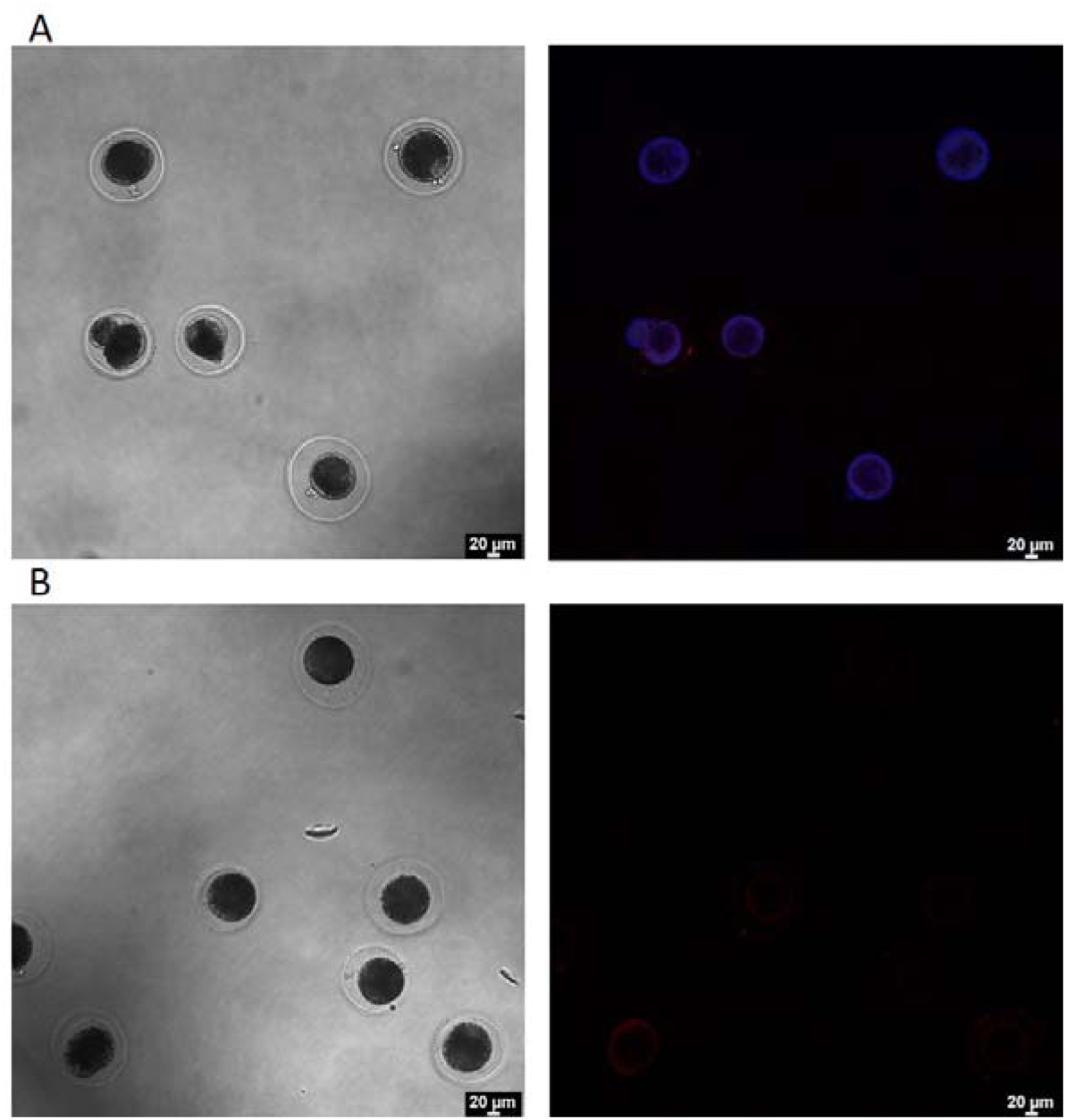
Electroporation of zona-weakened zygotes. Electroporation of pronase-treated zygotes with pooled FDs (3, 70, and 500 kDa) (A) and 2000 kDa FDs (B).

## Discussion

This series of studies investigated the impact of electroporation parameters on the delivery and fluorescence intensity of FDs in porcine zygotes and their subsequent development into blastocysts (Table 3). Our findings highlight the nuanced interplay between molecular size, electroporation conditions, and embryo viability, providing valuable insights for understanding and optimizing electroporation protocols in porcine zygotes.

The electroporation of 3 kDa, 70 kDa, and 500 kDa FDs into IVF zygotes allowed the calibration of fluorescence intensity settings to avoid crosstalk between channels. Despite the persistent red autofluorescence from the zona pellucida after FD electroporation in post-IVF zygotes, accurate fluorescence measurements were still attainable. We observed a significant and expected effect of FD size on fluorescence intensity, with smaller molecules exhibiting higher fluorescence values.

In the context of gene-editing, the delivery and functionality of different forms of Cas9 play a crucial role in editing efficiency. Compared to Cas9 mRNA, the use of RNP (ribonucleoprotein) complexes accelerates the gene editing process due to the shorter waiting time before genome editing occurs(4). However, the smaller combined size of the Cas9 protein (around 160 kDa) and sgRNA (around 34 kDa), compared to Cas9 mRNA (approximately 1,530 kDa) plus sgRNA, might also be a critical factor for increased editing efficiency. The concentration of Cas9 in zygotes has been shown to increase the likelihood of mutation events(13, 20, 21) emphasizing the importance of molecule size for efficient targeted mutations.

Polarity combination settings did not uniformly affect fluorescence across FD sizes, though polarity had a significant effect in the delivery of 3 kDa FDs, with reversed poring polarity increasing fluorescence values for this size-specific molecule. Reversing polarity improves transfection efficiency by addressing the suboptimal accumulation of molecules at the positive electrode (22). Although the transfer of molecules is theoretically influenced by the transfer pulses, it appears that this transfer predominantly occurs when the poring pulses are applied.

Electroporation of the 2000 kDa FD revealed that increasing the number of poring pulses beyond four did not enhance fluorescence intensity. This effect is likely due to the increased pulse rate failing to create larger pores in the embryo, which are necessary for the entry of large molecules. In contrast, Navarro et al. (2022) reported increased uptake with a higher pulse number, but their study involved a significantly smaller FD size (3 kDa tetramethylrhodamine)(23). Optimal results in our study were achieved at 4 pulses of 40 volts. Lower voltages were ineffective at delivering 2000 kDa FDs into zygotes, likely due to insufficient voltage to create adequate pores and the obstructive effect of the zona pellucida on large molecules. Increasing the voltage to 50 volts did not enhance fluorescence intensity compared to 40 volts and failed to target some of the zygotes. This is consistent with findings in bovine embryos, where increasing voltage beyond a certain threshold did not improve molecular permeability (9). Our results also demonstrated that higher voltages significantly affect blastocyst formation rates, a finding consistent with multiple studies on the electroporation of zygotes(9, 13, 24). These findings emphazise the importance of balancing voltage and pulse number to maximize transfection efficiency without compromising embryo viability.

Pre-IVF electroporation of zygotes successfully eliminated the red autofluorescence observed in post-IVF zygotes following FD delivery, regardless of the FD size. This phenomenon may be attributed to structural changes in the zona pellucida proteins that occur after fertilization. It is possible that FD particles become trapped in the zona pellucida of post-IVF embryos, leading to the observed autofluorescence. A statistical assessment between these pre- and post-IVF embryos was not possible due to the variability in ovary collection days. Literature data indicates that delivering macromolecules in mice is more challenging in zygotes than in mature oocytes, with zygotes blocking the entry of molecules larger than 110 kDa and oocytes blocking molecules larger than 170 kDa (25). Nevertheless, we successfully demonstrated the delivery of molecules up to 2000 kDa in both zygotes and oocytes.

The trend of increased fluorescence intensity with higher voltage levels in pre-IVF embryos was consistent with the results observed in post-IVF embryos. However, no significant differences were found across the analyzed groups. While this electroporation procedure appeared effective for molecule delivery, the blastocyst formation rate remained low at voltages higher than 20 V. This is likely due to the high voltage levels, as the 0 V control and the 20 V group did not show significant differences from the IVF control. Previous studies have achieved viable blastocyst formation rates following oocyte electroporation(15). The discrepancy in our study could be attributed to differences in electroporation devices or variations in the composition of maturation and fertilization media. Zona-weakened zygotes resulting from pronase treatment did not show improved 2000 kDA FD uptake at four pulses of 30 volts. Pronase treatment is used to increase the permeability in zygotes, increasing mutation rates and molecule entry in mice (12, 13). However, studies in pigs have indicated that mutation rates do not significantly differ between embryos with intact zona pellucida and zona-weakened embryos(26). In contrast, the same study reported that zona-free zygotes exhibit higher mutation rates compared to zona-intact embryos, albeit at the cost of meticulous isolation of individual zygotes in specialized wells and low blastocyst development rates.

In summary, while electroporation effectively delivers FDs of various sizes into porcine zygotes, the efficiency and viability are highly dependent on the precise calibration of electroporation parameters. The ultimate objective is to achieve successful electroporation of plasmids, which typically exceed 1000 kDa in size. While there are structural differences between plasmids, which are supercoiled, and dextrans, which have a globular structure, this study contributes valuable insights into correlating electroporation conditions with molecular size.

## Supporting information

Suppl. Mat.

## Acknowledgments

The authors gratefully acknowledge the financial support of the DFG (TRR 127), and a short-term stipend from the HGNI (TiHo, Hannover). This manuscript represents a partial fulfillment of a Ph.D. thesis of the first author (JPF) at the Graduate School of the Foundation of Veterinary Science in Hannover (TiHo), Germany.

## Author contributions

J.P.F. and W.K. designed the experiments and conducted the data analysis. P.H. and P.K. carried out oocyte retrieval, IVF, and parthenogenetic activation. J.P.F. and P.K. performed electroporation and analyzed the blastocyst formation rates. J.P.F. conducted microscopic and fluorescence measurements. All authors contributed to editing the manuscript

## Ethics Declaration

The authors declare no conflict of interest.

## Data Availability

Original Figures can be found in: https://zenodo.org/records/13358710?preview=1&token=eyJhbGciOiJIUzUxMiJ9.eyJpZCI6IjM2OTJlYTYyLWJiZDktNDUzYi04MWYwLWIzOGI3MzUwOGY5YSIsImRhdGEiOnt9LCJyYW5kb20iOiJjYjRjZTcxODUyZTU0ZWNmYmZlN2NmOWE2MzlmOWVkZCJ9.YbXrhHIYGa5uGfsfSZUh-YljRqmncLK9j-n_Py7khN7juPjw4-v3msa29-uf_NAScwOrYYB4DBt4X1Zd4K9cTw

