## Supplementary material for "Optimizing Electroporation Parameters for Efficient Delivery of Large Molecules into Pig Zygotes Using Fluorescent Dextrans from 3 to 2000 kDA": Suppl. Mat.

*Juan Pablo Fernández

- Institute of Farm Animal Genetics, Friedrich-Loeffler-Institut, Neustadt, Germany

2.Paul Kielau

- Institute of Farm Animal Genetics, Friedrich-Loeffler-Institut, Neustadt, Germany

3. Petra Hassel

- Institute of Farm Animal Genetics, Friedrich-Loeffler-Institut, Neustadt, Germany

6. Wilfried A. Kues

- Institute of Farm Animal Genetics, Friedrich-Loeffler-Institut, Neustadt, Germany,

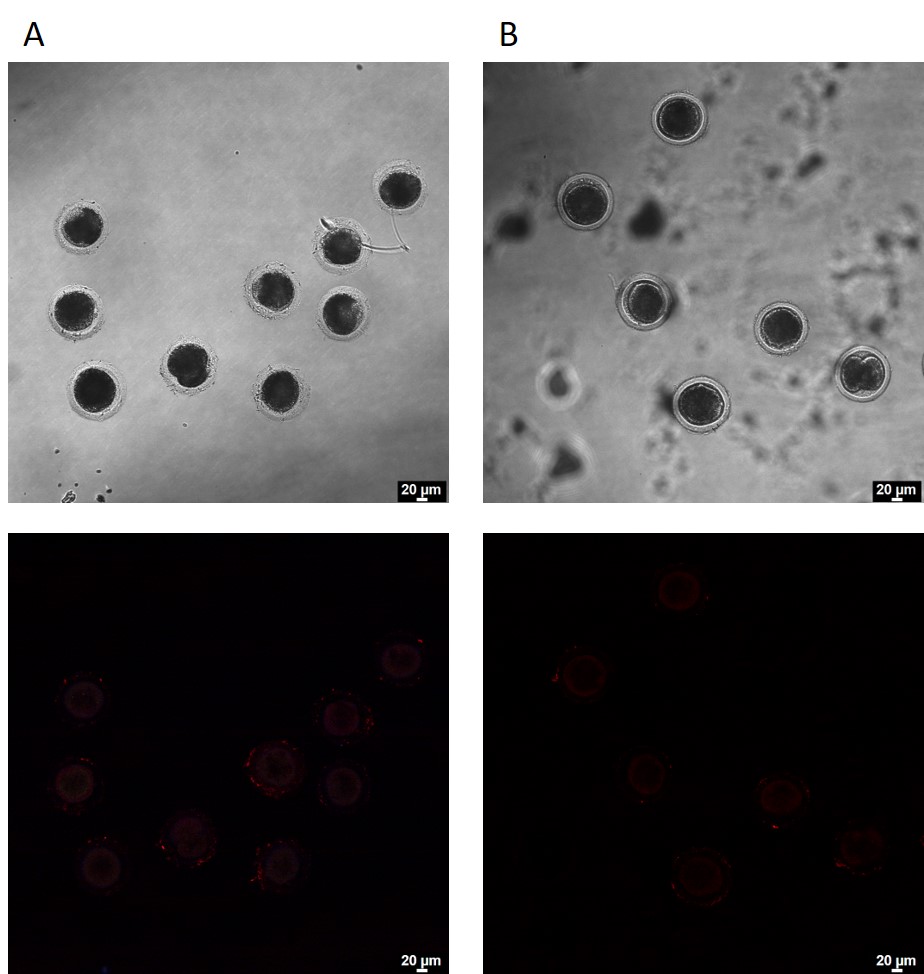

**Suppl. Figure 1:** Fluorescence recordings of IVF zygote electroporated at 30 V with 5 (A) and 6 (B) poring pulses
